# The *Helicobacter pylori* ribosomal silencing factor RsfS is required for low-growth states and chronic infection

**DOI:** 10.64898/2026.03.28.715003

**Authors:** Yasmine O. Elshenawi, Skander Hathroubi, Angela E. Lane, Megan Hetzel, Karen M. Ottemann

**Author notes:** SPARTHA Medical SAS Microbiology R&D Laboratory, CRBS 1 Rue Eugène Boeckel, 67000 Strasbourg, France.

## Abstract

*Helicobacter pylori* is a prevalent bacterial pathogen that chronically colonizes the human gastric epithelium, but the bacterium’s physiological mechanisms that promote this are understudied. Dormancy and low growth are known to facilitate other microbial chronic infections. A critical feature of low growth states is the down regulation of ribosome translational activity via regulation factors. The *H. pylori* genome is predicted to encode only one ribosome regulation factor, called RsfS (Ribosomal Silencing Factor S). In other bacterial species, RsfS prevents ribosome assembly by binding to a protein called L14 on the 50S large ribosomal subunit. Although *H. pylori* RsfS has not been experimentally investigated prior to this work, it conserves key residues, suggesting it is a bona fide RsfS homolog. To investigate phenotypes associated with *rsfS*, the gene was deleted and mutant phenotypes characterized. *H. pylori rsfS* null mutants had no defects during exponential phase but had viability defects in stationary phase and low growth factor conditions. Additionally, *rsfS* null mutants could not form biofilms, and instead were only able to form monolayers of multicellular aggregates. These defects were corrected by the re-introduction of *rsfS* in a second site on the chromosome. To explore whether *rsfS* is required *in vivo*, a mouse model was employed. *rsfS* mutants initially colonized in low numbers in both the glands and total stomach but were unable to develop robust long-term colonization. This work supports that *H. pylori* requires RsfS for survival in low growth states and to maintain chronic infections in the host.

**Importance:** *H. pylori* chronic infections are difficult to cure in part because *H. pylori* is proposed to adopt low-growth states known to render bacteria tolerant to antibiotics. One key signature of a low growth state includes low translation via ribosome regulation factors. Unlike other bacterial species, *H. pylori* contain only one known ribosome regulation factor called Ribosomal Silencing Factor S (RsfS). This gene was previously found to be transcriptionally upregulated in at least one low growth state, biofilms. In this work, we found that *H. pylori rsfS* is required for this microbe to thrive in low growth states and during infection. This study is one of only two studies that investigates the phenotypes of *rsfS* knockout mutants in any bacterial species and the first to address knowledge gaps in ribosomal regulation by *H. pylori in vivo*.

## Introduction

*Helicobacter pylori* is a gram-negative bacterium that colonizes the human stomach and establishes chronic infections. These infections can lead to gastrointestinal disorders such as peptic ulcers, gastric mucosal lymphoma, and gastric cancer (Chen et al., 2024; Kim et al., 2023; Malfertheiner et al., 2005). *H. pylori* is a globally prevalent pathogen that infects 40% of the population in developed countries and 80%-90% of the population in developing countries (Perez-Perez et al., 2004). In 1994, the World Health Organization classified *H. pylori* as a Class-1 carcinogen and the microbe has often made lists of priority pathogens due to the severe disease consequences (Wroblewski & Peek, 2013). *H. pylori* is often acquired in childhood, and is able to persist for the lifetime of the host (Czinn, 2005). *H. pylori* infections can be cured with antibiotics but generally require elevated dosages compared to other bacterial infections. For example, current recommendations suggest treatment for 2 weeks with 2-3 antibiotics combined with a proton pump inhibitor (Y.-C. Lee et al., 2022). These triple therapy treatments have been reported to fail 20-25% of patients treated in the United States (Schoenfeld, 2024; Vakil et al., 2004). The severity of *H. pylori* disease combined with its difficult treatment highlights a need for improved understanding of factors that feed into this challenge.

*H. pylori* forms chronic infections that last the life-time of a human host (Bruce & Maaroos, 2008). Many other bacterial pathogens also form chronic infections that occur in a variety of organs, including lungs (Cookson et al., 2018), central nervous system (Giovane & Lavender, 2018), gastrointestinal tract (Fantry et al., 2016), urinary tract (Hjelmager et al., 2025), and skin (Ki & Rotstein, 2008). One emerging idea is that these bacteria adopt low-growth states or dormant forms in the host to allow them to limit immune detection and thus facilitate chronic infections (Fisher et al., 2017; Rhen et al., 2003). These low-growth states are well known to contribute to treatment failures, largely because antibiotics mostly target processes that occur in growing cells, e.g. cell wall creation, translation, transcription (Da Silva et al., 2024; Rittershaus et al., 2013; Young et al., 2002). These low-growth chronic infection states have been well documented in *Mycobacterium tuberculosis* (Sakamoto, 2012), *Pseudomonas aeruginosa* (Garcia-Clemente et al., 2020), *Staphylococcus aureus* (Tuchscherr & Löffler, 2016)*, Salmonella* (Ehrhardt et al., 2023), and *Escherichia coli* (Hannan et al., 2012), but studies in *H. pylori* have lagged behind. There is some evidence, however, to suggest that *H. pylori* similarly adopt a low growth state during chronic infection. Gastric biopsy analysis from clinical patients have reported evidence of low-growth states due to the observation of morphological forms of *H. pylori* associated with dormancy (Reshetnyak & Reshetnyak, 2017a) and a growth mode associated with slow growth such as biofilms (Carron et al., 2006; Coticchia et al., 2006). *H. pylori* forms biofilms during *in vitro* conditions, and cells in this state are highly tolerant to antibiotics (Elshenawi et al., 2023; Windham et al., 2018). The fact that *H. pylori* can adopt these low-growth states and exhibits antibiotic tolerance has supported the idea that low-growth states might form *in vivo* and contribute to the difficulty of achieving complete eradication (Reshetnyak & Reshetnyak, 2017b).

Slow bacterial growth is a multifaceted phenotype in which multiple changes lead to low metabolic activity. One such change is ribosomal inactivation. Inactivated ribosomes have been previously associated with bacterial dormancy, older biofilm subpopulations, and states of low metabolic activity in various bacterial species (Chai et al., 2014; Kaprelyants et al., 1993; Y. Li et al., 2021; Williamson et al., 2012). Many microbes employ multiple ribosome-inactivating proteins. These include proteins that induce hibernation or silencing by promoting either ribosome dimerization or blocking ribosomal subunits association (Prossliner et al., 2018). *H. pylori*, in contrast, is predicted to encode only one ribosome-inactivating protein, a ribosomal silencing factor called Ribosomal Silencing Factor S (RsfS). RsfS (RsfA in other bacterial species) prevents the joining of 30S subunits to 50S subunits, and thus prevents the formation of active 70S ribosomes (Fig. 1A) (Fatkhullin et al., 2022). RsfS homologues are highly conserved across bacterial species as well as mitochondria (Häuser et al., 2012). RsfS protein homologues and their binding to the 50S subunit protein L14 have been structurally characterized in several bacteria, including *E. coli, S. aureus* and *M. tuberculosis* (Häuser et al., 2012; Khusainov et al., 2020; X. Li et al., 2016). Previous studies have also reported the conservation of RsfS homologues across bacterial and eukaryotic species and have confirmed binding between L14 and RsfS using the yeast two-hybrid method in *E. coli, T. palladium*, and the human mitochondrial homologue (Häuser et al., 2012). Overall, RsfS-type proteins are widespread across all domains of life and appear to conserve interactions with the ribosome.

**Fig. 1.**
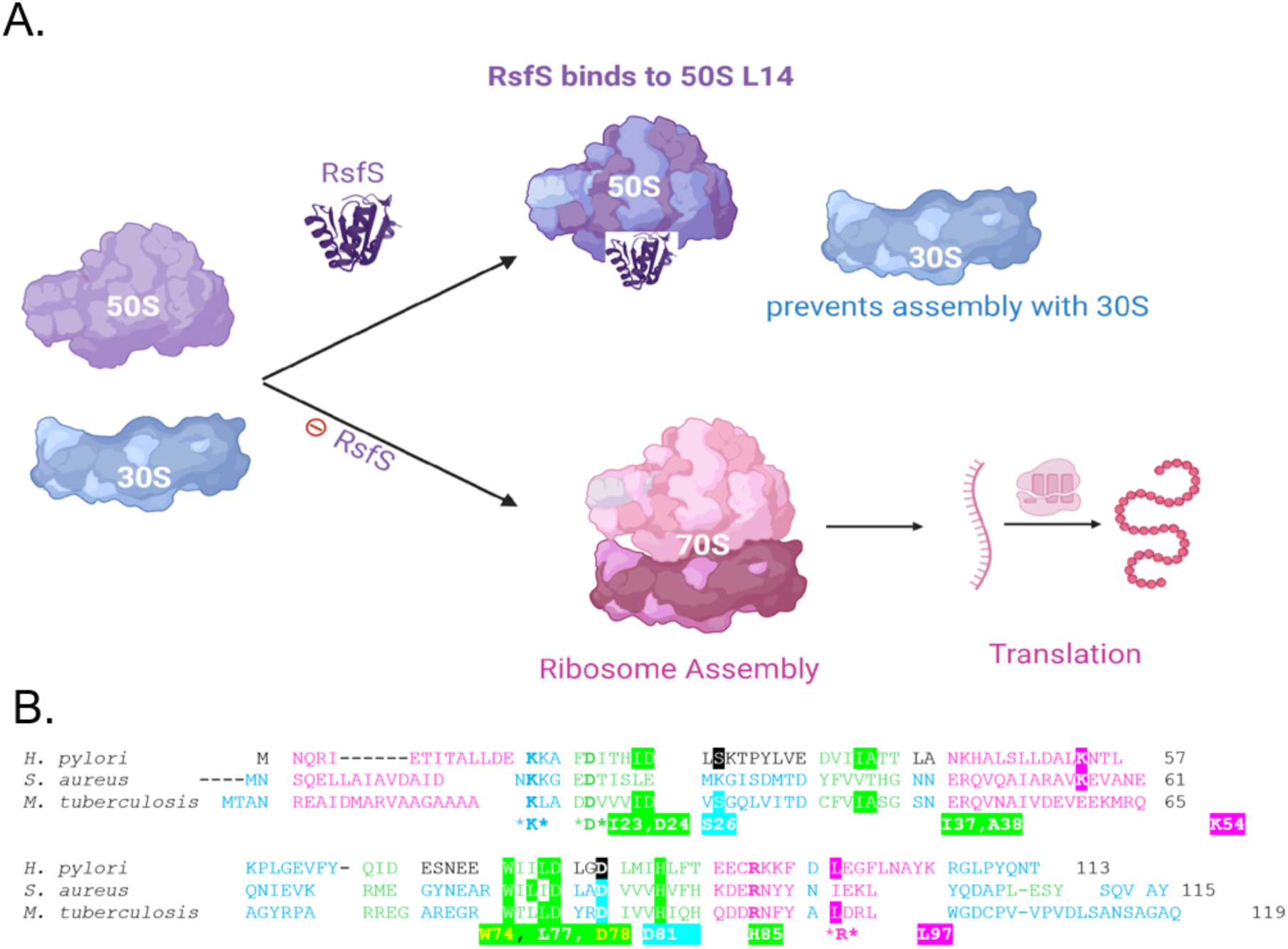
*H. pylori* contains RsfS Homologues. (A) Model for how the RsfS ribosomal silencing protein functions by binding to L14 protein on the large ribosomal subunit 50S, preventing assembly with the 30S subunit and formation of a 70S ribosome that can translate mRNA to polypeptides. **(B)** RsfS protein alignment for *H. pylori*, *S. aureus*. and *M. tuberculosis*. Secondary structures as predicted by alpha fold are color coded as the following: α-helices (pink), β-sheet (green) and coils (blue). RsfS residues that were found to be critical for binding to L14 via crystallography structures that are conserved in *H. pylori* are annotated in highlighted text beneath the sequence alignment. *Residues that were not identified in previous studies but conserved in all three species.

The only study that investigated the phenotypes associated with RsfS-type proteins was in *E. coli* (Häuser et al., 2012). *E. coli rsfA (*the *rsfS* homolog) null mutants had growth defects in stationary phase and in low-amino acid conditions, consistent with the idea that ribosomal silencing is important in these low-growth situations (Häuser et al., 2012). Additionally, the authors used a reporter gene assay and ribosome profiling to quantify assembled 70S versus separated 30S and 50S subunits. The findings led the researchers to suggest that *E. coli* RsfA prevents ribosome assembly under growth limiting conditions (Häuser et al., 2012). These findings established a premise for ribosome silencing mechanistically promoting bacterial survival during low growth states.

To date, no studies have examined the function of ribosome regulation in *H. pylori*. While *rsfS* transcript levels were reported to be elevated in biofilm populations compared to planktonic cells (Hathroubi et al., 2018). Ribosome populations and their regulators remain largely understudied in this bacterium. In this study, we investigate the conservation of RsfS and phenotypes associated with it in *H. pylori*. Our analysis confirms that *H. pylori* conserves critical residues in RsfS known to facilitate binding with L14, in agreement with previously reported protein-protein binding predictions (Humphreys et al., 2024). By creating *rsfS* knockout mutants, we determined that the loss of this protein resulted in growth defects under some conditions, including stationary phase, low nutrient conditions, and biofilm formation, but not others such as exponential phase and normal-nutrient media. These defective phenotypes could be corrected via genetic complementation with *rsfS* in a secondary site. In addition, *H. pylori* lacking *rsfS* mutants showed weak growth *in vivo.* These mutants initially colonized mouse stomachs in low numbers but were not able to maintain robust populations long term. Overall, our results indicate that *rsfS* expression plays a critical role in *H. pylori* low growth states *in vitro* and colonization *in vivo;* these phenotypes provide potential insights into how *H. pylori* regulate its ribosomal population to promote survival and sustain chronic and life-long infections.

## Results

### *H. pylori* RsfS conserves important residues

*H. pylori* is predicted to encode only one ribosome silencing factor, RsfS, annotated as HP1414 in the *H. pylori* 26695 (Tomb et al., 1997) or HPYLSS1_1340 in the SS1 genome (Draper et al., 2017). Because *H. pylori* RsfS has not been previously investigated, we first analyzed the protein sequence of *H. pylori* RsfS compared to homologues including those with validated binding interactions in which key residues were identified (Fatkhullin et al., 2022; Khusainov et al., 2020; X. Li et al., 2016). *H. pylori* RsfS encodes a 113 amino acid protein, predicted to fold into the same structure as characterized RsfS proteins. Amongst the bacterial species with validated RsfS ribosomal silencing (Fatkhullin et al., 2022; Khusainov et al., 2020; X. Li et al., 2016), the *H. pylori* RsfS was 21.5% percent identical (ID) to the homolog in *S. aureus* versus 19.3% percent ID to that in *M. tuberculosis*. Both *M. tuberculosis* and *S. aureus* contain somewhat longer forms of RsfS totaling 119 amino acids and 115 amino acids respectively. Previous studies have shown that particular residues in *S. aureus* and *M. tuberculosis* RsfS homologues drive protein-protein interactions with the ribosomal protein L14 (Fatkhullin et al., 2022; Khusainov et al., 2020; X. Li et al., 2016). Several of these *H. pylori* RsfS residues were conserved in *S. aureus* (Lys54, Asp81, Asp78, and Trp74) and *M. tuberculosis* (Ile23, Asp24, Trp74, Leu77, Asp78, His85, Ser26, Leu97) despite the low percent ID (Fig. 1B). Only three residues (K15, D19, R92) were found to be conserved across all three bacterial species that were not identified as critical binding residues (Fig. 1B). Thus is seems highly likely that *H. pylori* RsfS conserves the ability to interact with L14 based on this residue conservation, overall conserved secondary structure, and published bioinformatic analysis (Humphreys et al., 2024).

### *H. pylori* Δ*rsfS* mutants have growth defects compared to WT that are exacerbated in low nutrient conditions and with repeated disruptions

Next, we sought to analyze how the loss of *rsfS* would impact *H. pylori*. An *H. pylori* SS1 mutant lacking *rsfS* was created by replacing the full coding sequence of the *rsfS* gene with a chloramphenicol resistance marker (*cat*) to generate the strain SS1 Δ*rsfS*::*cat*, called Δ*rsfS* This mutation did not result in a polar effect on the downstream genes as evidenced by analysis of RNA levels using RT-qPCR (Supplemental Fig. 1). We then determined the growth phenotype of the resultant strain across a growth curve, starting in standard *H. pylori* rich media consisting of brucella broth with 10% heat-inactivated fetal bovine serum (FBS) (BB10). Using a 96-well microplate reader assay and measurements of optical density, the *H. pylori* Δ*rsfS* mutant had initially lower OD_600_ measurements compared to the WT strain, consistent with a somewhat enhanced lag phase (Fig. 2A). After ∼ 10 hours, a time point correlating to mid-exponential phase, the Δ*rsfS* mutant growth rate had increased, such that the mutant and WT strains had similar OD_600_ values at the late exponential time point, 20-hours. During the early and late stationary phases (24-36 hours), the OD_600_ values of the Δ*rsfS* strain remained steady, while the OD_600_ of WT increased to a modest degree (Fig. 2A). These results suggest that *H. pylori* can grow relatively well in rich media without *rsfS*.

**Fig. 2.**
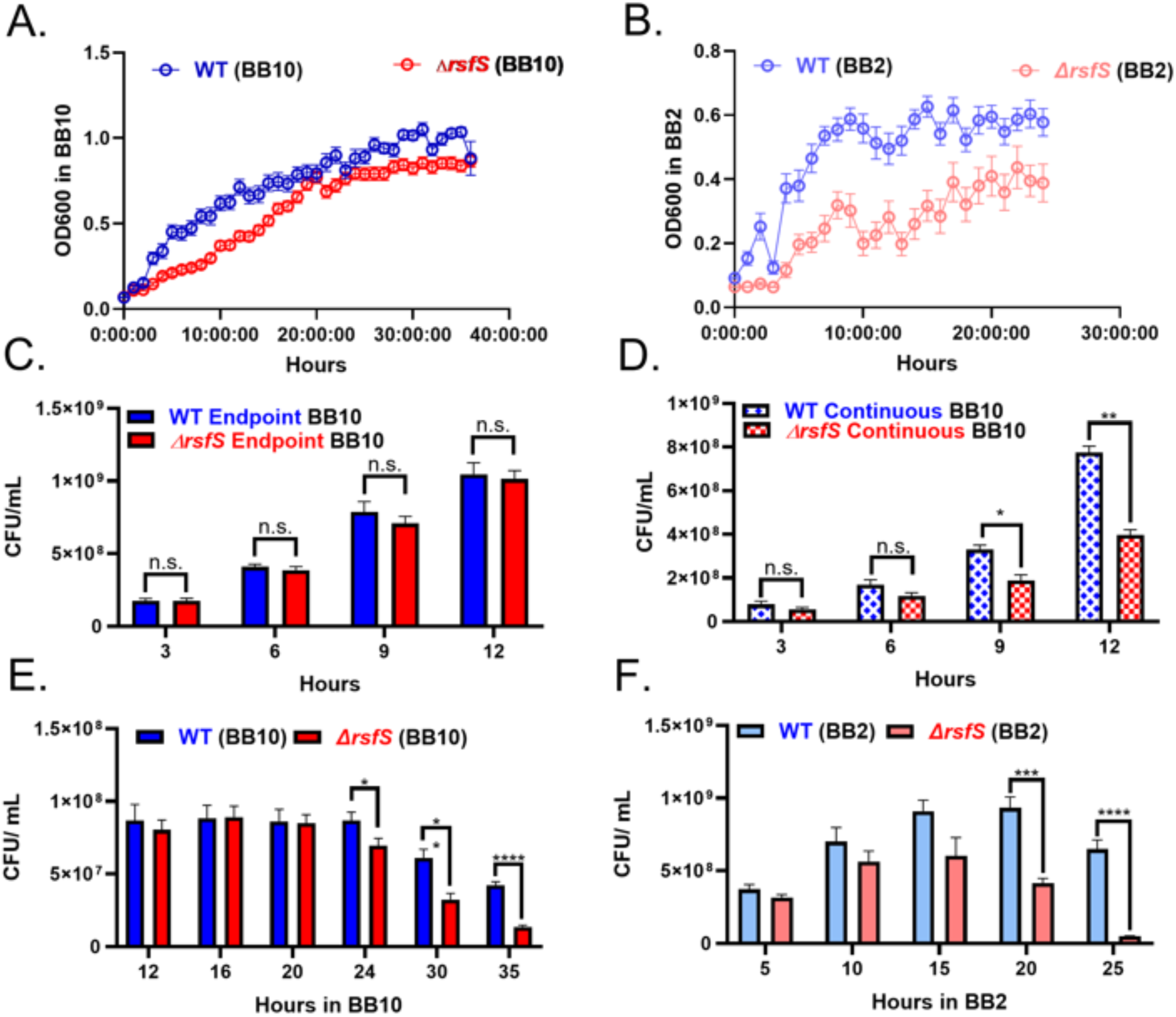
Growth analysis of *H. pylori* Δ*rsfS* mutant. Growth analysis of *H. pylori* SS1 WT (blue and light blue) and its isogenic *ΔrsfS* strains (red and light red) grown in either BB10 or BB2. **(A-B)** Growth analysis carried out in 96-well microtiter dishes. Biological replicates (n=4) and technical replicates for each timepoint (n=30-75). **(C-F).** Growth analysis carried out using flasks in the incubator (biological replicates n= 4-6) with measurements of colony-forming units at the indicated times. **(C)** Continuous growth approach in BB10. **(D-F)** Endpoint growth approach in **(D)** BB10 or **(E)** BB2. Pairwise T-test analysis was used to compare WT to mutant at matched times, with P values indicated as <0.001 (***) and <0.0001 (****).

Carrying out the same experiment but in low nutrient conditions (brucella broth with only 2% FBS, BB2) resulted in an enhanced defect of the *H. pylori* Δ*rsfS* strain (Fig. 2B). Both WT and the mutant strains achieved lower OD_600_ compared to those in BB10 and appeared to reach stationary phase sooner (Fig. 2B). The Δ*rsfS* strain, however, had consistently lower OD_600_ than the WT (Fig. 2B). These results suggest that loss of *rsfS* caused *H. pylori* to have enhanced defects in low serum conditions compared to high serum ones (Fig. 2B).

We next repeated the growth analysis but plated the bacterial samples to examine live bacteria. In this case, the bacteria were grown in tubes shaking in the incubator. A single culture was employed and repeatedly sampled, an approach referred to as continuous growth (Supplemental Fig. 2). Using this approach and rich medium (BB10), the Δ*rsfS* mutant produced significantly fewer CFU at the 9-hour time point relative to the WT control. This difference became more severe after 12 hours of incubation (Fig. 2C). We noted that this method had disrupted conditions at each time point, with more disruptions over time. This analysis suggested the *rsfS* mutant might be sensitive to changes in conditions encountered during disruptions associated with sampling.

To evaluate whether the sampling regime was affecting the *rsfS* mutant, we sought a second growth approach that would eliminate repeated sampling and disruptions to conditions. This outcome was accomplished using an endpoint assay, in which the entirety of an individual sample was used and repeated disruption was eliminated (Supplemental Fig. 2). Using the same timepoints as the continuous growth assay (Fig. 2C), no differences in CFU were observed between WT and two distinct Δ*rsfS* mutants (Fig. 2D, Supplemental Fig. 3). The endpoint approach further revealed that the *H. pylori* Δ*rsfS* mutant did not have a defect compared to WT in the first 20 hours of growth, exponential phase (Fig. 2E). However, as the cells transitioned into stationary phase, loss of *rsfS* created *H. pylori* with significant growth defects that became progressively worse as the cells progressed through stationary phase (Fig. 2E).

Continuing the endpoint analysis approach, we repeated the growth analysis of the Δ*rsfS* mutant in BB2. In BB2, the Δ*rsfS* mutant produced less CFU/mL than the WT control across all timepoints, but the difference between the Δ*rsfS* strain and WT did not become statistically significant until stationary phase time points at 20-25h (Fig. 2F). Similar results were observed with an independent Δ*rsfS* mutant (Supplemental Fig. 3). Compiled, these results suggest that *rsfS* is required for *H. pylori* to withstand repeated disruptions in growth conditions, for growth in low serum media, and for survival in stationary phase.

### *H. pylori* Δ*rsfS* mutants are weak biofilm formers

Previous work showed that *H. pylori rsfS* expression was significantly elevated in biofilms compared to free-floating cells (Hathroubi et al., 2018). It was not known, however whether RsfS was important for *H. pylori* biofilm growth. *H. pylori* SS1 produces robust biofilms under static, microaerobic conditions in BB2 media after three days, before dispersing at four days (Hathroubi et al., 2018). We thus quantified biofilm formation of the *H. pylori* Δ*rsfS* mutant under these conditions, using two distinct *rsfS* mutant alleles. Δ*rsfS* mutants formed significantly less biofilm than the WT strain (Fig. 3A, Supplemental Fig. 3). Specifically, at the 3-day timepoint the mutant had ∼3-fold less biofilm as measured by crystal violet staining. To evaluate whether the ΔrsfS strain might have delayed biofilm, the samples were incubated for an additional day, but the mutants still displayed approximately 2-fold less biofilm than the WT strain (Fig. 3A). These results suggest that *rsfS* is important for *H. pylori* biofilm growth.

**Fig. 3.**
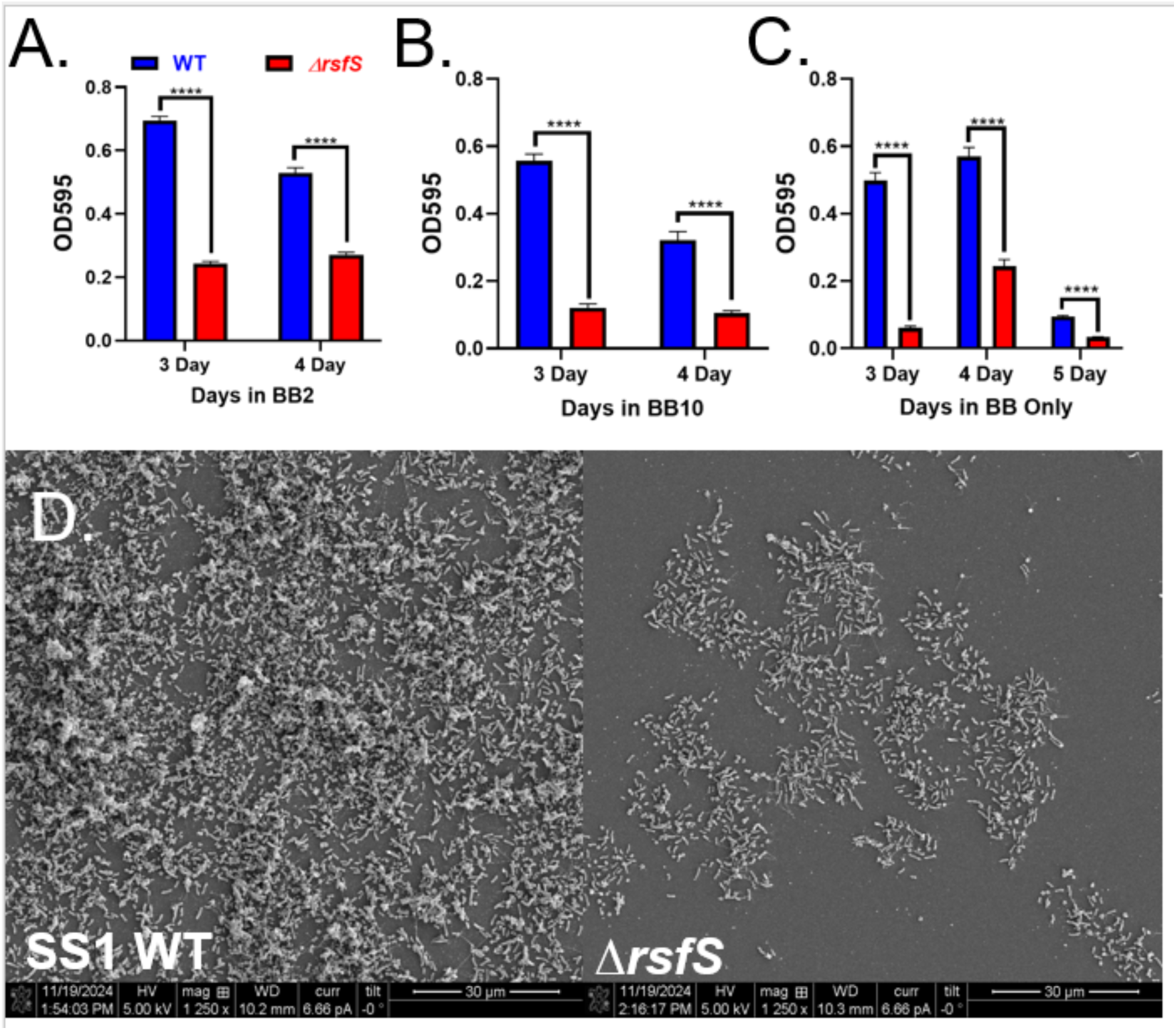
*H. pylori* Δ*rsfS* mutants are weak biofilm formers. Temporal analysis of biofilm formation in various media. Four biological replicates of the WT control (blue) and *ΔrsfS* (red) biofilms were grown in 96-well plates, stained with crystal violet and OD_595_ was measured at the indicated times. **(A)** BB2 (technical replicates 105-252). **(B)** BB10 (technical replicates 35-115). **(C)** No serum (BB only) (technical replicates 38-136). Statistical analysis comparing WT and mutant using T test. **(D)** Biofilms in BB2 were formed on glass cover slips for 3 days and imaged using scanning electron microscopy (SEM) for both *H. pylori* WT and Δ*rsfS.* Pairwise T-test analysis was used to compare WT to mutant, with P values indicated as <0.0001 (****).

We wondered whether the Δ*rsfS* mutant would retain this significant biofilm defect when grown in alternative media. We hypothesized that the Δ*rsfS* mutant BB2 growth deficit (Fig. 2) might be driving the biofilm defect, so we explored media with higher serum concentrations where the Δ*rsfS* mutant has no exponential phase defect (Fig. 2). However, even with 10% serum (BB10), the *ΔrsfS* strain retained its biofilm defect (Fig. 3B). Specifically, the mutant formed 4.6-fold less biofilm than the WT at 3 days (Fig. 3B). In addition to BB10, we also explored whether the *rsfS* mutant would display a biofilm defect in media with no serum, based on a report with *S. aureus* defining FBS as a potential biofilm inhibitor (Abraham & Jefferson, 2010). At 3 days, the SS1 WT control produced about 8-fold the amount of biofilm as the *ΔrsfS* strain, a statistically significant difference that reduced to 2-fold at 4 days prior to dispersal at 5 days (Fig. 3C). Taken together, these results provide strong evidence that the *H. pylori ΔrsfS* strains produced significantly less biofilms relative to the WT across all time points and media conditions.

To gain further insight into the biofilm defect associated with loss of *rsfS*, we used scanning electron microscopy (SEM) to visualize *H. pylori* WT and *ΔrsfS* strain biofilms grown for 3-days in BB2. The SS1 WT strain produced robust networks of multilayer aggregates and flagellar filaments, as reported previously (Hathroubi et al., 2018), whereas the Δ*rsfS* mutant strain had fewer aggregate structures, and aggregated cells generally only formed monolayers (Fig. 3D). These findings supported the crystal violet assays, strongly indicating that the *rsfS* is essential for biofilm formation in *H. pylori*.

### The Δ*rsfS* mutant phenotypes can be complemented by exogenous expression of *rsfS* in an intergenic locus but not *rdxA*

The above results suggest that *H. pylori* RsfS is critical for biofilm formation, growth in low nutrient and changing conditions, and stationary phase survival (Fig. 2, 3). To confirm these roles of *rsfS*, we sought to complement the mutant by inserting a copy of *rsfS* into a heterologous locus. We initially used the *rdxA* gene, a chromosomal locus commonly used for complementation in *H. pylori* due to its classification as a neutral locus (Chan et al., 2015; Damke et al., 2019; Howitt et al., 2011; Kwon et al., 2000; Liechti & Goldberg, 2013, 2013; Liu et al., 2022; McGee et al., 2005; Nguyen et al., 2025; Rader et al., 2007; Rocha et al., 2025; Smeets et al., 2000; Sycuro et al., 2010; Terry et al., 2005; G. Wang et al., 2012; X. Wang et al., 2025; Zimmerman et al., 2024). We created two *rdxA::rsfS* alleles and used them to transform both WT and the Δ*rsfS* mutant (Supplemental Fig. 4). Exogenous expression of *rsfS* via the *rdxA* locus, however, did not complement the Δ*rsfS* planktonic growth or biofilm defects (Supplemental Fig. 5). This led us to question the neutrality of loss of *rdxA* and examined *rdxA* knockout mutants for growth defects. Surprisingly, loss of *rdxA* alone caused defects in both late stationary phase and biofilms (Supplemental Fig. 5). This phenotype is being explored in a separate manuscript, but this outcome suggested that *rdxA* was not a suitable locus for complementing these types of phenotypes.

As an alternative to *rdxA*, we designed a construct that would place a copy of *rsfS* into an intergenic region that has been used previously (Langford et al., 2006) (Supplementary Fig. 4). As with the *rdxA* integrated constructs, a copy of *rsfS* with 200 base pairs upstream was integrated between the coding sequences for HP0203 – HP0204. After verifying the correct integration, we examined whether the defects observed in the Δ*rsfS* knockout mutants would be corrected. Both biofilm formation and stationary phase defects of the Δ*rsfS* mutant were corrected by placement of a copy of *rsfS* in the intergenic region (Fig. 4). This outcome strongly suggests that these defects were due to the loss of *rsfS*.

**Fig. 4.**
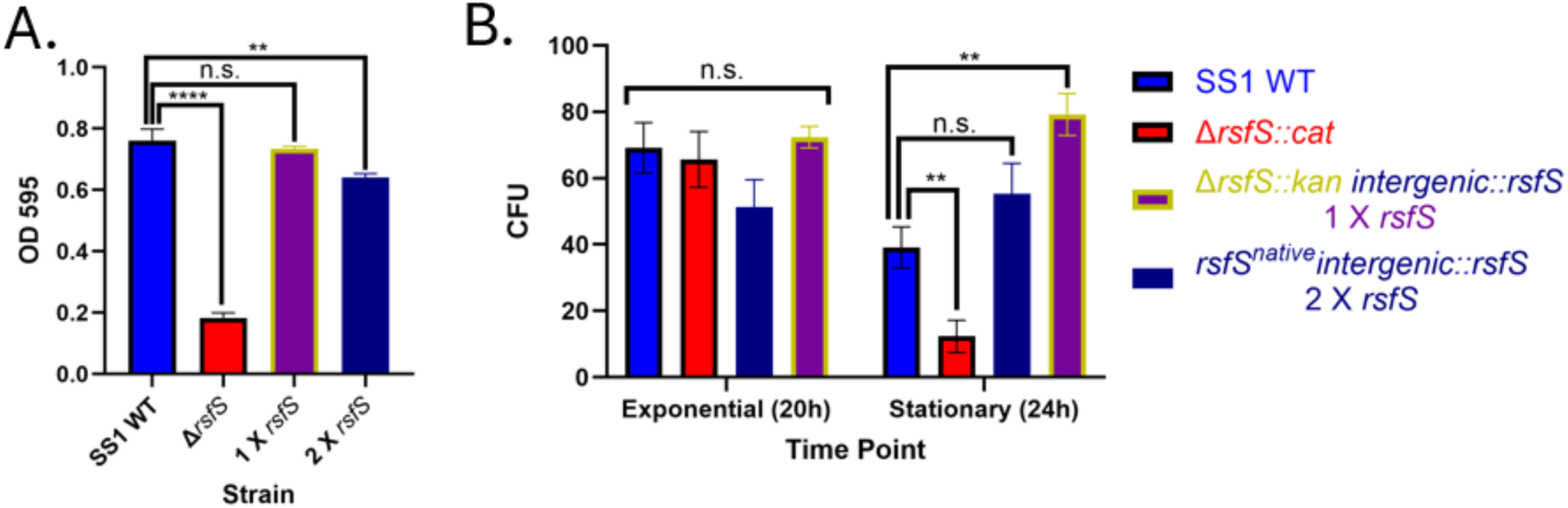
*rsfS* mutants are complemented by insertion of an exogenous copy of *rsfS*. *rsfS* with 200 bp upstream and a chloramphenicol resistance marker (*cat*) was integrated into the intergenic region between HP0203 and HP0204 in either the Δ*rsfS* mutant (1X *rsfS*) or WT (called 2X *rsfS).* **(A)** Biofilms were grown in BB2 for the 3 days and analyzed by crystal violet staining (OD_595_). 4 biological replicates per strain, with technical replicates of WT, 20; *ΔrsfS::cat*, 21; 1X *rsfS*, 84; and 2X *rsfS,* 100. Students T test analysis was conducted comparing each strain to WT, p value <0.0001, ****. **(B)** The endpoint growth approach was used to compare CFU yields during exponential (t=20h) and stationary (t= 24h) phase growth in BB10. T-test analysis was used to compare each strain to WT (P-value <0.01, **).

### *rsfS* possesses an upstream promoter that contributes to its expression

Because the *rsfS* complementation constructs also included an upstream 200 base pairs and the putative promoter, we evaluated expression of *rsfS* in the context of these various non-native locus constructs. Previous work demonstrated that the *rsfS* is the first gene in a four gene operon (Supplemental Fig. 1A), with a transcriptional start site mapped in *H. pylori* strain 26695, at an A 26 base pairs upstream of the *rsfS* ATG (Bischler et al., 2015). Two constructs possessed this promoter region as part of the 200 bp *rsfS* upstream section and no other known promoter. The construct that placed *rsfS* in the opposite orientation as *rdxA* (Δ*rdxA*::*rsfS* (1p)), and the construct that placed *rsfS* in the intergenic region (*IG::rsfS)* (Supplemental Fig. 4). After making these strains, transcripts levels of *rsfS* were examined using q-RT-PCR, normalized to the *gapB* reference gene to calculate ΔCq (Fig. 5). Strains bearing only the 200 bp upstream of *rsfS* expressed *rsfS* to levels that were within 1 cycle of the WT strain (Fig. 5A), and when normalized to *gapB* (Fig. 5B), resulted in ΔCq values that had a difference of ∼1.4 from the WT (Fig. 5C). These findings support that the *rsfS* promoter is indeed directly upstream of *rsfS*.

**Fig. 5.**
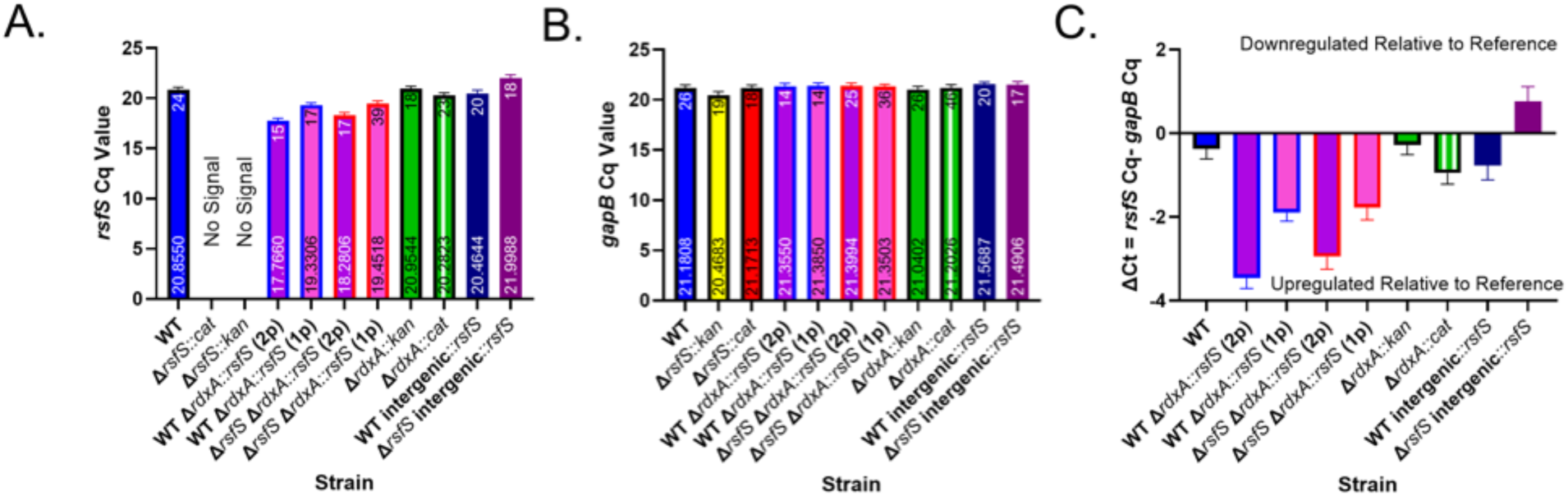
*rsfS* possesses an upstream promoter that contributes to its expression. qRTPCR was conducted on RNA isolated form 4-6 biological replicates to determine the expression **(A)** target *rsfS* gene **(B)** reference *gapB* gene to assure RNA quality. Cq Values are plotted with mean and standard deviations values of ∼0.3 to ensure qPCR and RNA quality within strains. Technical replicates are annotated at the top of the bars whereas means are annotated at the bottom on the bars. **(C)** ΔCt was calculated by subtracting the target gene Cq value (*rsfS*) by the reference gene Cq value (*gapB*).

### Overexpression of *rsfS* leads to elevated stationary phase survival

In addition to complementing the *rsfS* mutant, we also created strains that possessed two copies of *rsfS*, the native WT copy plus the complementing copy. Expression of *rsfS* in these backgrounds was analyzed as above by q-RT-PCR. With the intergenic *rsfS* construct, we noticed that while the second copy was expressed, combining it with the native *rsfS* did not produce higher expression than the single native copy alone (Fig. 5A). Indeed, having both copies actually resulted in lower expression compared to just one at either the native or intergenic locus.

Strains in which there was an additional *rdxA* promoter (Δ*rdxA*::*rsfS* (2p)) expressed the most *rsfS* transcripts, with about a ∼ 2-3 fold increase in *rsfS* expression relative to the WT (Fig. 5C). This *rsfS* double copy strain—with elevated *rsfS*--had only a slight effect on biofilm formation, significantly less biofilm than the WT parent strain but more than the control Δ*rdxA::kan* and Δ*rdxA::cat* strains (Supplement Fig. 5D). Overexpression of *rsfS* using the intergenic locus (Supplemental Fig. 4, Fig 5A&C) had a more substantial effect on stationary phase CFU, leading to close to 40% more CFU compared to WT (Fig. 4B). This outcome suggests that *rsfS* expressions have an important effect on controlling stationary phase CFU.

### The *rsfS* gene is needed to sustain long-term infections

Our results above support that *rsfS* plays an important role in modulating *H. pylori* growth *in vitro* in specific conditions. To evaluate how these phenotypes would translate to stomach colonization, the *in vivo* behavior of the *H. pylori rsfS* mutant was evaluated. Mice were orally infected with equal amounts of either *H. pylori* WT or *rsfS* mutant, using strains that had additionally been transformed with the GFP-expressing plasmid pTM115 (Fig. 6A) (Keilberg et al., 2016). The mice were left infected for one or four weeks. After that time, stomachs were collected and divided into the two distinct anatomical regions called corpus and antrum. Colonization levels were assessed by plating for CFU and gland populations were analyzed by microscopy with the Bacterial Localization in Glands (BLIG) approach (Keilberg et al., 2016). After one week, both strains were detectable in the glands in both regions of the stomach, as assessed by percent of glands infected and the number of *H. pylori* per gland (Fig. 6B-D). In both regions and by both measures, the WT was found in higher numbers, with significant differences identified in the number of bacteria per gland and in total CFU (6C-E). By four weeks, the difference in the number of glands occupied had also become statistically significant in both regions (Fig 6B). In all measures, the amount of the Δ*rsfS* mutant strain decreased during this period, while the WT mostly increased. Of note, we were unable to recover live Δ*rsfS* mutant bacteria at four weeks, even though we could detect some GFP+ strains. In summary, the *in vivo* results indicate that lacking the *rsfS* gene causes *H. pylori* to have poor ability to initiate and maintain infections.

**Fig. 6.**
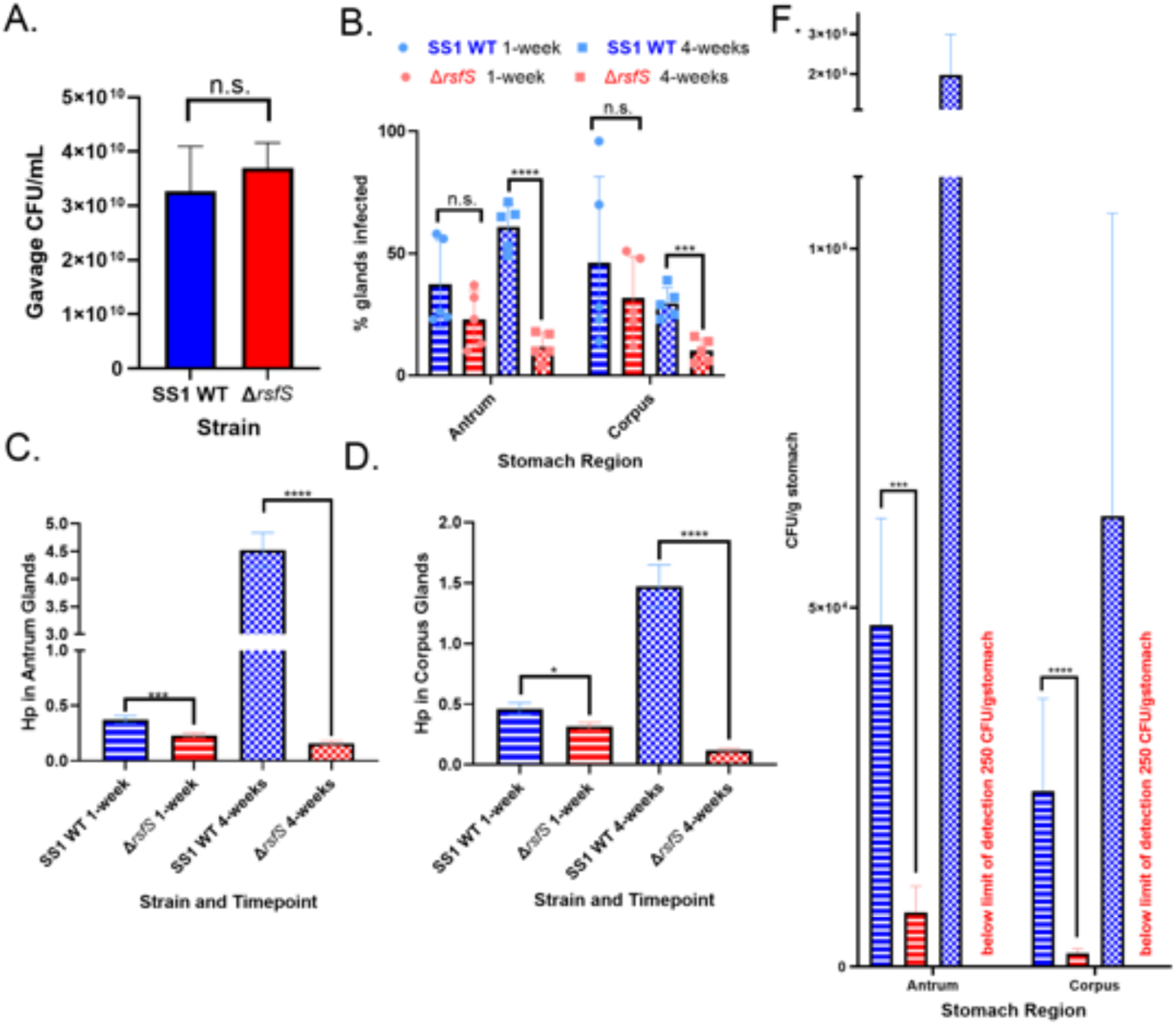
Δ*rsfS* mutant colonizes mice initially but fails to sustain long-term infections. Two sections of the stomach (antrum and corpus) from n=20 mice were used to compare the phenotypes of *ΔrsfS* and the control SS1. **(A)** Six technical replicates were serially diluted and plated to assure similar amounts of inoculums were used. Using BILG, 100 gastric glands (both occupied and unoccupied) from each stomach section were analyzed to calculate **(B)** the percentage of glands infected by *H. pylori,* the population of *H. pylori* per gland in the **(C)** antrum and **(D)** corpus. **(E)** Stomach tissue was then homogenized to culture live *H. pylori*. Statistical analysis was done using t-test p<0.1(*), p<0.001 (***), and p<0.0001 (****).

## Discussion

We report here that *H*. *pylori* requires its predicted ribosomal silencing factor, RsfS, for survival and growth in conditions that are associated with low-growth and *in vivo*. Slow growth or dormant states have been proposed to facilitate chronic infections (Bode et al., 1993; Reshetnyak & Reshetnyak, 2017a) and poor responses to antibiotic treatment in several microbes (Kaprelyants et al., 1993). A common feature in bacterial low growth states is low translation activity (Chai et al., 2014). Ribosome inactivation has been investigated in other species and determined to be an essential physiological mechanism for low growth states (Kaprelyants et al., 1993; Song & Wood, 2020; Williamson et al., 2012). Before this work, how *H. pylori* might use ribosome inactivation remained largely unstudied. Our analysis supports that the *H. pylori* RsfS conserves certain key residues with other well-studied RsfS homologs that are important for its interactions with the ribosomal L14 protein. Deleting *rsfS* from *H. pylori* decreased the microbe’s survival in stationary phase, low serum conditions, and biofilm assays, phenotypes that could be complemented by placement of an *rsfS* gene in an intergenic locus. This complementation also provided support for the prediction that there is a promoter upstream of *rsfS*. Additionally, deleting *rsfS* also caused severe deficits in *H. pylori’s* ability to initially colonize mice and impaired the ability of *H. pylori* to achieve the same levels of bacteria after four weeks as WT did. The combined findings in this study lead us to propose the model that *rsfS* is needed for *H. pylori* to survive challenging growth conditions and promote low-growth states that are critical in situations such as the stomach and biofilms.

RsfS operates by binding to the L14 protein on the 50S subunit and preventing the 50S and 30S subunits from assembling into a fully assembled 70S ribosome (Fig. 1A). The interaction of RsfS and L14 is highly conserved in multiple species, as assayed using a yeast two hybrid assay (Häuser et al., 2012; Parrish et al., 2007), and in a couple species using either crystallography or sucrose gradient methods (Fatkhullin et al., 2022; Häuser et al., 2012; Khusainov et al., 2020; X. Li et al., 2016). Our analysis here shows that the *H. pylori* RsfS protein conserves at least 12 residues that have been found in other RsfS homologs to be critical for L14 interactions (Fig. 1B). Additionally, the *C. jejuni* RsfS and L14 homologs have been shown to interact as part of a global yeast two hybrid screen (Parrish et al., 2007), thus supporting that RsfS/L14 binding may be conserved in species closely related to *H. pylori*. Consistent with our analysis, Humphreys *et al.,* 2024 reported the RsfS-L14 interaction is predicted to be conserved in *H. pylori.* as well as many other bacteria including *M. tuberculosis, S. aureus, Listeria monocytogenes* and *P. aeruginosa* (Humphreys et al., 2024). That work also suggests that RsfS proteins can interact with other proteins beyond L14. Specifically, they found that *P. aeruginosa* RsfS bound a PhoH-like protein called YbeZ (Humphreys et al., 2024), although the function of this interaction is not known. Thus, it is highly likely that *H. pylori* RsfS interacts with L14 but possibly has additional targets.

Although genetic knockout and complementation experiments are a common approach to investigate the role of genes in bacteria (Tong et al., 2023), only one study has applied this approach to study *rsfS* or the *E. coli rsfA* (Häuser et al., 2012). In that work, *E. coli ΔrsfA* had defects in stationary phase and nutrient conditions that could only be corrected by complementing with the entire four gene *rsfA* operon but not *rsfA* alone (Häuser et al., 2012). *E. coli rsfA* and *H. pylori rsfS* do not have the same operon structure. Replacing *H. pylori rsfS* with *cat* did not significantly affect the transcription of other genes in the operon, indicating our mutation was not polar as was observed in *E. coli* (Supplemental Fig. 1), and we were able to complement the mutant with only rsfS in an exogenous locus (Fig. 4).

The first phenotypes we investigated with the *H. pylori rsfS* null mutant were planktonic growth (Fig. 2). Three conditions were initially found to significantly delay the growth of *rsfS* knockout mutants: frequent disruptions in shaking for measurements obtained in a plate reader (Fig. 2A), low serum conditions (Fig. 2B), and repeated exposure to atmospheric conditions using a continuous growth curve approach (Fig. 2C). The growth impairment due to fluctuations in growth conditions matches the conclusions made with *E. coli rsfA* (Häuser et al., 2012), prompting these authors to conclude that RsfA played a role in adaptation to stressful growth conditions. When bacteria are starved for nutrients, they are able to mount a coordinated response that alters transcription, decreased ribosome biogenesis, and other effects (Akiyama & Kim, 2023). One known pathway is the stringent response that depends on a signaling molecule called ppGpp (Brown et al., 2016). Although ppGpp has not been extensively characterized in *H. pylori,* it has been associated with response to fluctuating growth conditions including nutrient starvation, stationary phase and aerobic shock (Mouery et al., 2006; Wells & Gaynor, 2006). Given that our findings support that *rsfS* is needed by *H. pylori* to mitigate stressful growth conditions, RsfS may be a part of *H. pylori’s* stringent response as well. Using the endpoint growth curve approach (Supplemental Fig. 2), we were able to mitigate these limiting growth factors that could complicate our analysis of the *rsfS* null mutant phenotypes (Fig. 2C). Without these additional complexities, it became clear that *H. pylori rsfS* knockout mutants have defects in stationary phase and low serum conditions, similar to phenotypes reported in *E. coli rsfA* mutants. Overall, these data further indicate that *H. pylori* RsfS homologues have a function that is conserved in *E. coli* toward responding to challenging growth conditions.

We report here that *H. pylori rsfS* is required for biofilm formation. Based on our endpoint growth curve experiment, we were able to identify a time point at which the *rsfS* mutant did not have a growth defect compared to WT (Fig. 2D). We used cultures grown to this time point, 20 hours, for biofilm and mouse inoculation to mitigate growth effects in these assays. Previous work had shown that *rsfS* was upregulated in *H. pylori* biofilms cells relative to unadhered cells (Hathroubi et al., 2018). In the work herein, we found that *H. pylori rsfS* mutants had consistent biofilm defects under a variety of conditions that could not be compensated for with longer incubations (Fig. 3), but could be complemented by exogenous *rsfS* expression (Fig. 4). SEM imaging further corroborated that the *rsfS* mutants were unable to form robust multi-layered aggregates as seen in WT. Given that biofilms also contain slow-growing populations, the findings appear consistent with the planktonic growth phenotype in noting that *H. pylori* needs *rsfS* during slow growth. Whether RsfS homologs are needed for biofilm formation in other bacteria remains to be determined.

Our first approach to complement the Δ*rsfS* mutant was to utilize the *rdxA* locus. This approach failed as, to our surprise, *rdxA* itself is needed for biofilm formation and stationary phase survival (Supplemental Fig. 5). Defective phenotypes that are similar between loss of *rdxA* and *rsfS* suggest that both are needed for challenging growth conditions. We therefore generated a distinct complementing strain, with *rsfS* inserted in a previously used intergenic region (Langford et al., 2006). This approach allowed the *rsfS* mutant phenotypes to be complemented, confirming *rsfS* itself is required for biofilm formation and stationary phase growth (Fig. 4). Overall, these results provide strong support for the idea that the *H. pylori* SS1 mutants described here are defective only for *rsfS*.

Previous work had predicted that *rsfS* was the first gene in a four gene operon (Supplemental Fig. 1A), with a transcriptional start site located 26 basepairs from the first *rsfS* codon, and a presumed promoter about 60 basepairs from the first codon (Bischler et al., 2015). q-RTPCR in this work confirmed that the 200 bp upstream of *rsfS* was sufficient for expression in both the *rdxA* and intergenic loci (Fig. 5). Furthermore, *rsfS* was still expressed when the *rsfS* gene and its upstream region were inverted within the *rdxA* locus (Fig. 5) further supporting the existence of a promoter in this 200 bp region.

To further investigate the role of *rsfS* in *H. pylori* growth *in vivo,* mice were infected with the mutant or WT and analyzed for both CFU and stomach gland populations. By some measures, the mutant was able to initiate a reasonable infection, e.g. the percentage of glands infected at 1-week were similar between the *rsfS* mutant and the WT (Fig. 6B), but at this time the *rsfS* mutant population in the glands and total CFU were significantly less than with WT (Fig. 6D, E). Thus, it can be concluded that the *ΔrsfS* mutant can establish initial colonization but at significantly lower numbers. After four weeks, the *ΔrsfS* mutant could be visualized in the glands (Fig. 6 B,C,D), but the bacteria were not able to be re-cultured from the mice (Fig. 6E). These results suggest that RsfS contributes to *H. pylori* colonization and persistence *in vivo*, consistent with the idea that this protein supports survival in low-growth states and under stressful conditions encountered in the mammalian stomach.

Our study finds that the only known ribosome regulation factor in *H. pylori,* RsfS is required for this bacterium to survive during slow growth and other challenging conditions including murine stomachs. Based on similarity to other well characterized homologues, *H. pylori* RsfS likely interacts with ribosomal L14 and contributes to ribosome silencing. How this process contributes to survival during slow growth is not well understood, with the idea that RsfS slows translation and/or protects ribosomes (Fatkhullin et al., 2022). Regardless, our work suggests that eliminating or blocking RsfS would render *H. pylori* less able to survive low-growth states and chronic infections *in vivo*.

## Supporting information

All supplemental figures

## Acknowledgements

The described project was supported by the National Institute of Allergy and Infectious Disease (NIAID) grants R01AI164682 and RO1AI116946 to K.M.O. The authors are grateful to Ottemann lab members Jashwin Sagoo for help with the transcriptional start site mapping, Dr. Jiunn Fong for help with cloning, Dr. Shuai Hu for help with mouse experiments and cloning approaches, Dr. Frances Yap for input on ribosomal regulation topics, Dr. Fitnat Yildiz for insight on biofilm phenotypes and all members of the Ottemann lab for support and input on this project.

## Methods and Materials

### Sequence Analysis for *H. pylori* L14 and RsfS

Clustal Omega program (https://www.ebi.ac.uk/jdispatcher/msa/clustalo) was used to generate guide trees that compared the genetic conservation of *rsfS* and *L14* of *H. pylori* relative to species that were investigated in Häuser, Roman, et al. (2012). Protein sequences were obtained via UCSC microbes genome browser (https://toolkit.engineering.ucsc.edu/website-support/maintenance/) and Open Reading Frames sequences using default parameters were obtained for analysis. The Clustal Omega program was also used to align RsfS and L14 protein sequences of *E. coli* (Grenier et al., 2014)*, S. aureus* (Kuroda et al., 2001)and *M. tuberculosis* (Lew et al., 2011) to *H. pylori* one by one to identify whether amino acid residues that were previously identified to be critical to RsfS-L14 binding are conserved.

### *H. pylori* Strains and Growth Conditions

*H. pylori* strain SS1 (A. Lee et al., 1997) was used as a wild type control for all assays and the parent strain to generate mutant strains (Table 1). All *H. pylori* strains were incubated at 37°C under microaerobic conditions with 5% O_2_, 10% CO_2_, and 85% N_2_. All chemicals are from Thermo Fisher or Gold Biotech unless otherwise indicated. Media consisted of Columbia Horse Blood Agar base with the addition of 0.2%-*β* cyclodextrin dissolved in dimethyl sulfoxide, 10µg of vancomycin per ml, 0.025 µg of cefsulodin per ml, 0.0125 U of polymyxin B per ml, 50µg of cycloheximide ml, 5% (vol/vol) defibrinated horse blood (Hemostat Labs). For antibiotic resistance marker selection, CHBA was supplemented with 15μg of chloramphenicol (Cm) per ml, 15μg of kanamycin (Kan) per ml or 1 μg of erythromycin (Erm) per ml. Liquid cultures were grown in Brucella Broth (Fisher Scientific) with 10% heat-inactivated Fetal Bovine Serum (Gibco) resulting in the media abbreviation BB10. For long term storage, strains were stored at −80°C in brain heart infusion media supplemented with 10% heat-inactivated fetal bovine serum, 1% (wt/vol) β-cyclodextrin, 25% glycerol, and 5% dimethyl sulfoxide.

**Table 1.**
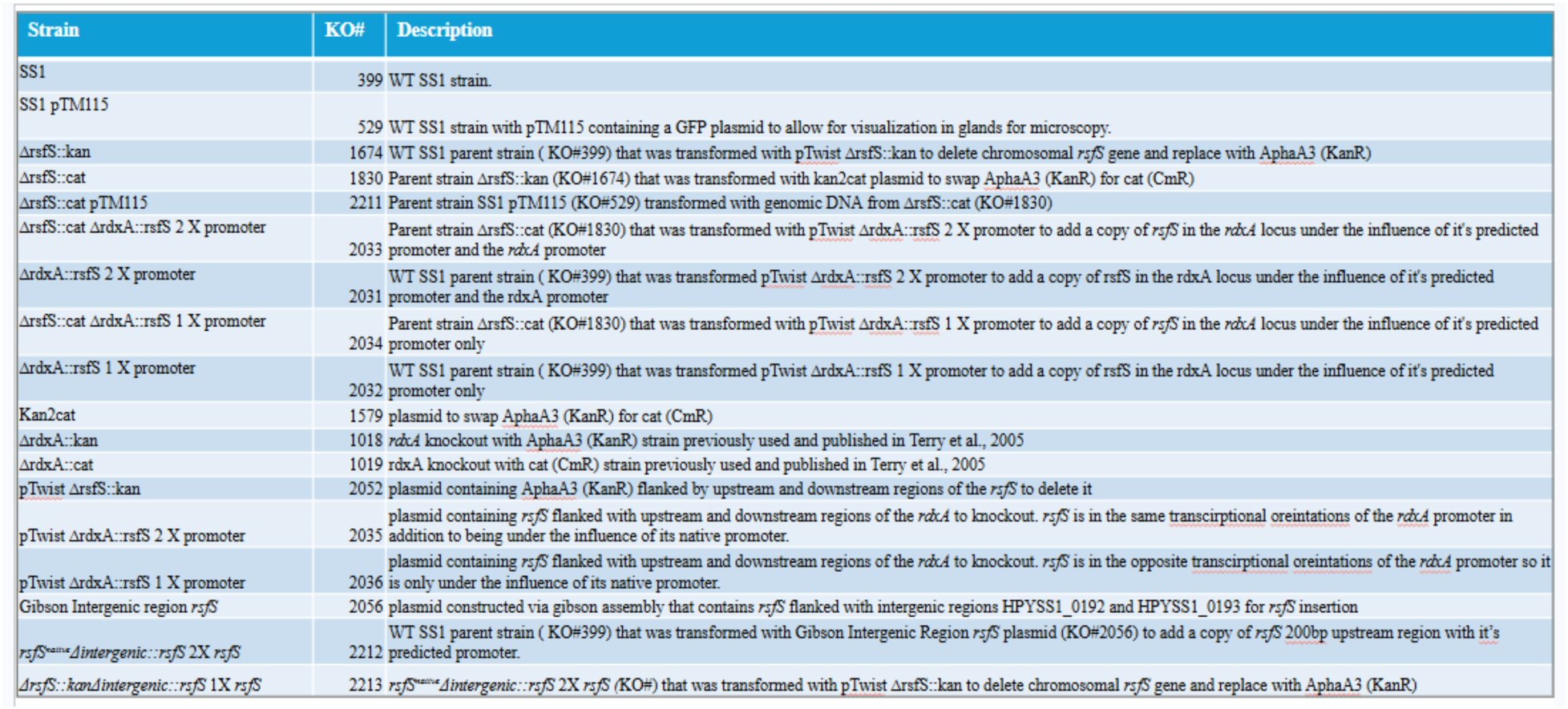
Strains and plasmids used in this study.

### Creation of *H. pylori* Mutants

All *H. pylori* transformations were done using natural transformation protocols, by either adding DNA directly to plate grown *H. pylori* or to *H. pylori* scraped from the plate into BB10 (Hu et al., 2023). In all cases, *H. pylori* was incubated on selective media until single colonies were detectable, and then single colonies were colony purified before collecting for storage at −80C. In all cases, genome modifications were analyzed using PCR products from genomic DNA preparations (Promega Wizard Prep) that flanked or were within the region or interest with Q5 High Fidelity 2X Master Mix. PCR products were purified using GFX Gel Band and PCR Purification Kit (Ge28-9034-71) analyzed by both agarose gel electrophoresis and DNA sequencing (Azenta), and in some cases by q-RT-PCR for gene expression.

To create *H. pylori* Δ*rsfS* deletion mutants (HP1414, HPYLSS1_1340 in the *H. pylori* 26695 (Tomb et al., 1997) and SS1 genomes (Draper et al., 2017) respectively, a synthetic construct called pTwist *ΔrsfS::kan* (Supplemental Fig. 2), was created that contains a kanamycin resistance gene (*aphA*3, Kan^R^) flanked by upstream and downstream regions of *rsfS,* designed to replace the *rsfS* gene (HPYSS1_1340/HP_1414) with the a*phA3* to generate *H. pylori ΔrsfS::kan.* This plasmid was stored in *E. coli* DH5α using ampicillin selection, and isolated using a miniprep approach with QIAprep Spin Miniprep Kit. The resulting plasmid was used for natural transformation to transform *H. pylori* SS1 WT to Kan^R^. To swap the *aphA* gene for chloramphenicol resistance, the *H. pylori* SS1 *ΔrsfS::kan* strain was transformed with a kan2cat plasmid (KO#1579, Table 1) containing a *cat* (Cm^R^) gene flanked by *aphA3* sequences and selecting for chloramphenicol resistance resulting in *ΔrsfS::<u>cat</u>* <u>(KO#1830)</u>. Genomic DNA (2µg) from *ΔrsfS::cat* was then used to transform into SS1 pTM115 (KO#529) by adding directly to patches of colonies that previously incubated for 5 hours. Transformations were incubated overnight and then streak plated onto CHBA supplement with Cm to select for colonies containing the *ΔrsfS::cat* allele. Transformants were then visualized under the microscope to check for GFP positive fluorescence to ensure the strain *ΔrsfS::cat* pTM115 (KO#). Transformants were analyzed by PCR using primers that flank the *rsfS* sequence, specifically targeting 263 bp upstream and 335 bp downstream relative to the *rsfS* sequence called *rsfS* flanking Forward and *rsfS* flanking Reverse (Table 2).

**Table 2.**
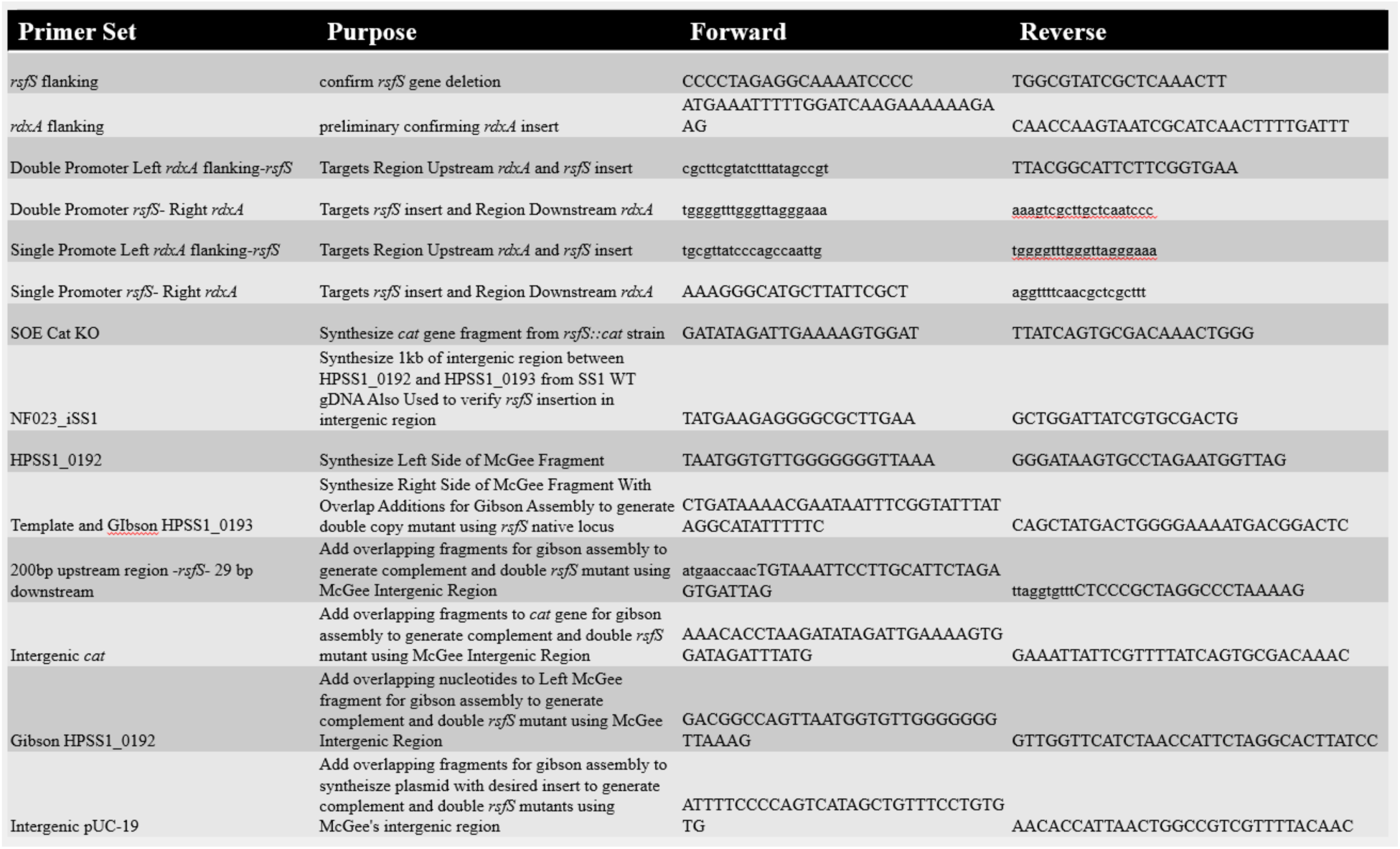
Primers used in this study.

To create complementation strains with *rsfS* insertions in the *rdxA* locus, synthetic plasmids (Twist Bioscience) were designed that contained the full *rsfS* gene and its potential promoter according to the annotations in the 26695 genome, an erythromycin resistant gene (*erm*) and sequences from *rdxA* to promote integration at that chromosomal locus (Supplemental Fig. 4). Plasmids were maintained in *E. coli* DH5α using ampicillin. *H. pylori* SS1 WT or *ΔrsfS* strains were used as parent strains for transformation and colonies were selected on CHBA supplemented with Erm to select for *rsfS*-*erm* integration in *rdxA*. Chromosomal integration of the *rdxA::rsfS* transformants was confirmed by PCR with primers (1) targeting the upstream region of the *rdxA* locus and the gene using primers “Double Promoter Left *rdxA* flanking-*rsfS”* and “Single Promoter Left *rdxA* flanking-*rsfS”* (Table 2); (2) targeting the *rsfS* gene from the insert and the downstream region of the *rdxA* locus using primers “Double Promoter *rsfS*-Right *rdxA”* and “Single Promoter *rsfS*-Right *rdxA”*.

Complementation strains were created by inserting *rsfS* in an intergenic region between HP_0203and HP_0203, as used in Langford et al. 2006. Gibson assembly was conducted using the following four fragments: (1) Left intergenic, (2) 200bp upstream region, *rsfS* ORF, 29bp downstream intergenic, (3) *cat* resistance cassette and (4) Right intergenic. The template containing *rsfS* with predicted promoter was obtained from *H. pylori* SS1 genomic DNA by PCR with primers *rsfS* flanking Forward and Reverse (Table 2), and then further amplified to synthesize the second fragment with overhangs for Gibson assembly. The second fragment was synthesized to include the *rsfS* ORF, 200bp upstream the *rsfS* ORF to account for a promoter and 29bp of intergenic region downstream *rsfS* ORF using the Gibson *rsfS* primer set (Table 2). The flanking intergenic regions were synthesized from SS1 WT gDNA using primer set NF023_iSS1 primer set and used as a base template to synthesize first and fourth fragments with Gibson overhangs. The first fragment was synthesized using the primer set Gibson 193 which also added overhanging sequences for Gibson assembly (Table 2). The template for the fourth fragment was initially generated from the purified intergenic region PCR product using primer set HPSS1_0192 (Table 2). The resulting PCR product was purified so Gibson overhangs could be added using primer set Gibson HPSS1_0192 creating the fourth fragment (Table 2). The third fragment containing the *cat* gene was then generated using purified PCR products generated from Δ*rsfS::cat* gDNA using *rsfS* flanking primer set (Table 2). The purified PCR product from *rsfS* flanking primers were then used to synthesize the *cat* gene template using primer set CAT SOE (Lowenthal et al., 2009). Gibson overhangs were then added to the *cat* gene template with primer set Gibson CAT primer set creating the third fragment (Table 2). All four fragments were synthesized and purified and stored in 4 degrees for no less than 24 hours prior to Gibson assembly with the plasmid backbone.

To construct the backbone to complete the Gibson assembly, pUC-19 plasmids were isolated from KO#137 (SY327 pUC19) using Qiagen miniprep kit (CAT# 27106). The pUC-19 plasmid was then linearized using High Fidelity Restriction Enzyme Hind III (CAT#R3104S) using manufacturer instructions, then immediately purified and used as template to add Gibson overhangs using Gibson Int-pUC-19. The resulting PCR product was purified immediately resulting in a plasmid backbone with Gibson overhangs added. The initial templates generated through PCR from Hp gDNA, the resulting fragments that were synthesized with Gibson overhangs and the linearized pUC-19 plasmid with Gibson overhangs were purified using Genomic DNA Clean & Concentrator™ Kits, Zymo Research CAT# 77001-150 (KT).

All PCR reactions for gibson assembly were conducted with Q5 Master Mix, NEB, Q5 2X Master Mix CAT# M0492S. Gibson assembly was then conducted using NEBuilder HiFi DNA Assembly Master Mix CAT#E2621L, fragment concentrations were determined by using the NEB HiFi calculator provided and used to make 20 uL reactions of 4 fragments and linearized pUC-19 plasmid with Gibson overhangs combined with 10uL of master mix. The reaction was incubated according to protocol recommendations for 4-6 fragment reactions, 50°C for 1 hour. The resulting plasmid was then transformed into *E. coli* DH5α and colonies that up took the Gibson plasmid were selected by supplement LB agar with Chloramphenicol (20µg/mL). Chloramphenicol resistant colonies were streak plated and plasmids were isolated using the QIAprep Spin Miniprep Kit were sent to AZENTA for whole genome sequencing. Upon the verification of the insert sequence remaining intact without SNPs, mutations or truncations, the resulting DH5α isolate containing the *pIG::rsfS* (KO#2056) was cultured in liquid LB supplemented with Cm and mixed with 80% glycerol and stored in −80°C long term.

*H. pylori* transformations were then conducted with the plasmid *pIG::rsfS* (KO#2056) isolated via QIAprep Spin Miniprep Kit to insert *rsfS* gene copy in the defined intergenic region (Langford et al., 2006). *H. pylori* strain SS1 WT (A. Lee et al., 1997) was used as a parent strain with transformation of *pIG::rsfS* (KO#2056) plasmid on CHBA supplemented with Cm and confirmed by PCR using primer set NF023_iSS1 primer set (Table 1, 2). To create the complementation construct, this strain was transformed with pTwist *ΔrsfS::kan* (KO# 2052) to delete the native copy of *rsfS*; the resulting transformation was selected on CHBA supplemented with Kan and Cm and verified by PCR using primer set *rsfS* flanking (Table 1, 2).

### *H. pylori* growth assays

For growth assays in the plate reader, *H. pylori* strains were grown initially overnight in BB10 with shaking for 12-15 hours in tubes or flasks. Overnight cultures were normalized to OD_600_ of 0.1 and used to inoculate 96-well flat bottom, non-treated clear polystyrene with low evaporation lid (Corning cat#7200656)to a final volume of 200uL in the desired media (BB10 or BB2). Microplate growth assays were conducted using CLARIOstar^Plus^ Microplate Reader (BMG LABTECH) with microaerobic conditions (double orbital shaking 200 rpm) with 5% O_2_, 10% CO_2_ and 85% N_2_ at 37°C. OD_600_ was measured hourly throughout the assays.

Overnight BB10 cultures were inoculated from CHBA-plate grown *H. pylori* and incubated for 12-15 hours in BB10, using 3 mL in a 15 mL falcon-style tube. Endpoint and monoculture replicates were normalized to an OD_600_ of 0.1 at the start of the experiment, followed by incubation (shaking) under microaerobic conditions with 5% O_2_, 10% CO_2_ and 85% N_2_ at 37°C. Samples for optical density (OD_600_) and colony forming units (CFU/mL) were collected by removing 100uLfor each type of analysis at the desired time point.

### Biofilm Assays

Biofilm phenotypes were analyzed using a protocol previously adapted to *H. pylori* (Hathroubi et al., 2018). *H. pylori* strains were grown for 20h with shaking in BB10 using 3 mL in 15 mL falcon-style types. After growth, the cultures were diluted to an OD_600_ of 0.15 using fresh media (BB only, BB2 or BB10). 200uL per well was inoculated in sterile 96-well polystyrene microtiter plates (Costar#3370). Plates were incubated in static microaerobic conditions (5% O_2_, 10% CO_2_, 85% N_2_) for 72h. Media was removed from the wells via pipetting, and the remaining biofilms were stained with 300uL of crystal violet (0.1% wt/vol) at room temperature for 3 minutes. The crystal violet solution was then removed by pipetting and the wells were washed with 300ul of 1X phosphate-buffered saline (PBS) twice. After these steps, the plates were dried for 20 minutes at room temperature and 300uL of ethanol (95% vol/vol) was added to the wells and incubated at room temperature for 10 minutes. After this period, optical density at 595 nm was measured.

To collect biofilms for SEM imaging, the initial overnight growth and back dilution were the same except the back diluted cultures were inoculated into µ-Slide 8-well glass bottom plates (ibidi, Germany #80826) followed by the placement of 12mm diameter 1/2oz EM cover slips (Electron Microscopy Sciences) in each well diagonal, so biofilms could form on the slide at the liquid-air interface on the acute angle side. The cover slips were static in microaerobic conditions as described above for 72h. The cover slips were then removed and rinsed with PBS before fixation at room temperature for 1h in 2.5% glutaraldehyde. After fixation, serial dehydration steps were conducted by placing the coverslips for 10 minutes, room temperature in progressively higher amounts of ethanol: 25%, 50%, 75%, 90% and 100% (100% step was repeated twice). The coverslips containing the fixed biofilms were then left in the final 100% ethanol solution until SEM imaging was conducted shortly afterwards within the hour.

### RNA Extractions

RNA extractions done using the Purelink RNA Minikit (CAT#12183018A) from H. pylori grown as described above. Bacterial cells were collected, suspended in TRIzol, and stored in −80°C until RNA isolation. TRIzol-suspended samples were thawed at room temperature before the addition of chloroform. Samples were shaken by hand until thoroughly mixed prior to being incubated at room temperature for 3 minutes. Samples were centrifuged at 15,000 × g for 15 minutes at 4°C which caused phase separation of the mixture into a lower pink phenol-chloroform phase, an opaque white interphase, and a clear upper phase containing RNA. The phase containing RNA was transferred to a new tube and an equal volume of 70% ethanol (final concentration ∼35%) was added and mixed by vortexing and vigorous inversion to disperse any precipitate. The homogenized mixture was placed in a Purelink RNA Minikit spin cartridge, and manufacturer’s instructions followed with an additional on-column DNase treatment using Turbo DNA-Free Kit (Invitrogen ThermoFisher CAT# AM1907) for 15 minutes at room temperature. After this step, the column was washed four times at 4°C with wash buffer 1 and then wash buffer 2, followed by a repeat of both steps. After washing, the column was dried by centrifugation at 15,000 × g for 1 minute at 4°C. RNase–Free water was added to the membrane and incubated at room temperature for 1 minute prior to eluting the extracted RNA by centrifuging for 1 minute at 15,000 x g at 4°C. RNA yield and quality was quantified using a Nanodrop spectrometer. RNA was stored at either −80°C long-term or in4°C for immediate processing into cDNA.

### cDNA synthesis

RNA was synthesized into cDNA using High-Capacity cDNA Reverse Transcription Kit with RNase Inhibitor (Applied Biosystems/Thermo Fisher CAT#4374966) using the manufacturer’s recommendations. 1ug of RNA for each sample was used for cDNA synthesis and diluted with RNase-Free water to meet the total recommended volume of 10ul. One technical replicate of cDNA synthesis was conducted per biological RNA replicate. A no RNA template control (NTC) and a no-reverse transcriptase control (No RT) were added to assess contamination.

### RT-qPCR protocol

RT-qPCR was performed using reagents from the Power SYBR™ Green PCR Master Mix kit (Applied Biosystems#4367659) and a Bio-Rad CFX96 Real Time PCR Detection System. All reactions were carried out with 1µL of the cDNA reaction (synthesized from 1µg of RNA) mixed with 5 µM each of the forward and reverse primers for the target gene, 10 µl of PCR Master Mix, and water for a total volume of 20 µL per reaction a. The thermal cycling conditions were executed as the following: Step#1-95°C for 10 minutes, 40 cycles of denaturing at 95°C for 15 seconds and Annealing/extension at 60°C for 1 minute with a final melting curve step for 5 seconds. No template and no-reverse transcriptase controls were used as controls. Four to five technical replicates for each primer set were conducted for each biological RNA replicate.

### Soft Agar Assays

Soft agar plates that were made using Brucella broth (BB) media, 2.5% heat-inactivated Fetal Bovine Serum and 0.35% (w/v) agar and supplemented with *H. pylori* selective antibiotics: 50µg of cycloheximide per mL, 10µg of vancomycin per ml, 5µg of cefsulodin per ml, 2.5 U of polymyxin B per ml (Thermo Fisher or Gold Biotech). Soft agar plates were inoculated by using a sterile micropipette tip and picking bacteria from CHBA plates and inserting the tip containing bacteria into the soft agar. The soft agar plates were incubated in microaerobic conditions at 37°C for 6 days prior to measurement.

### Ethics Statement

The University of California Santa Cruz Institutional Animal Care and Use Committee approved all animal protocols and experiments (protocol OTTEK2405). All animal procedures used were in strict accordance with the Guide for the Care and Use of Laboratory Animals. Female C57BL/6N mice (Helicobacter free; Charles River), ages 4-6 weeks, were housed at the University of California Santa Cruz animal facility

### Mouse Infections

Prior to infection, *H. pylori* strains were grown overnight for 16-20h hours in BB10, collected by centrifugation, normalized to an optical density at 600 nm of ∼ 1, and then 500 µl was used via gavage. Infection cultures from *H. pylori* SS1 pTM115 or its isogenic Δ*rsfS*::*cat* derivative were serially diluted and plated to assure similar amounts of CFU between both strains were used in the infectious dose. Mice were sacrificed at 1 and 4 weeks. using CO_2_ narcosis followed by cervical dislocation to confirm death prior to dissection to extract the stomach for both infection methods.

### Stomach Extraction

The stomach was extracted by severing the stomach-esophageal junction and the antrum-duodenum sphincter using scissors. The stomach was then opened along the lesser curvature from antrum to for stomach and then the contents of the stomach (food and debris) were removed by rinsing the internal side of the stomach in 1X phosphate-buffered saline (PBS). The stomach was then spread out using pins and the forestomach was removed. The stomach was sectioned into corpus and antrum based on the anatomical orientation of the stomach and tissue coloration, with the corpus identified as pink versus the antrum as translucent/white.

### Stomach Bacterial Numbers

The tissue collected for total bacteria number was weighed then homogenized using the Bullet Blender (Next Advance) with 1.0-mm zirconium silicate beads. The resulting homogenate was serially diluted and plated on CHBA supplemented with 10ug/ml nalidixic acid, 20ug/ml bacitracin and 1000X HP (5 µg/ml trimethoprim and 8 µg/ml amphotericin B) to select for *H. pylori*.

### Gland isolation

The gland isolation protocol utilized in this study was a protocol from a previous study (Keilberg et al., 2016). Dissected gastric tissue from both regions was cut into approximately 6 X 4 mm rectangles, which were then incubated at 4°C for 3-4 hours with slight shaking in Dulbecco’s phosphate-buffered saline (DPBS) supplemented with 5 mM EDTA. Tissue samples were transferred to DPBS supplemented with 1% sucrose and 1.5% sorbitol then subsequently stained with 2uL of Hoechst DNA stain (Life Technologies). The tissue suspensions were put on ice to settle. A total of 100 glands were counted for gavage infection and 200 glands were counted for micro-pipette infection. Bacteria per gland (sample size for gavage: 100 glands) were counted in each region to calculate how many glands were infected and the average bacteria per gland.

