## Supplementary material for "The *Helicobacter pylori* ribosomal silencing factor RsfS is required for low-growth states and chronic infection": All supplemental figures

A.

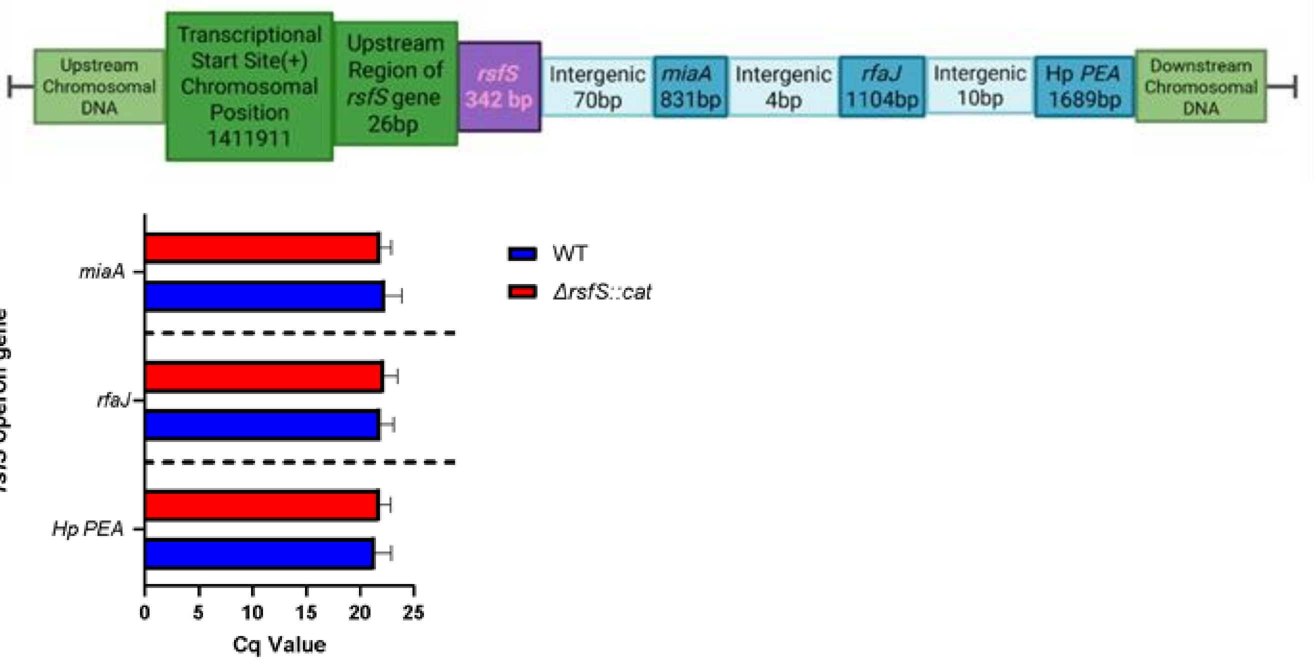


**Supplemental Figure 1. *rsfS* operon and downstream gene expression in Δ*rsfS* mutants. (1)** The *rsfS* operon contains a total of four genes in following order: *rsfS* (HPYSS1_1340), *miaA* (HPYSS1_1341), *rfaG* (HPSS1_1342) and *Hp PEA* (HPYSS1_1343). **(2)** Transcriptional expression of *rsfS* operon genes in Δ*rsfS::cat* mutant strain suggest the *rsfS* deletion/insertion is not a polar mutation. RNA was extracted from plate grown bacteria. Cq values between Δ*rsfS::cat* and SS1 WT control produce a standard deviation of approximately 0.3 which is indicative that there is no differential transcription of the operon genes due to a polar effect caused by the *rsfS* deletion.

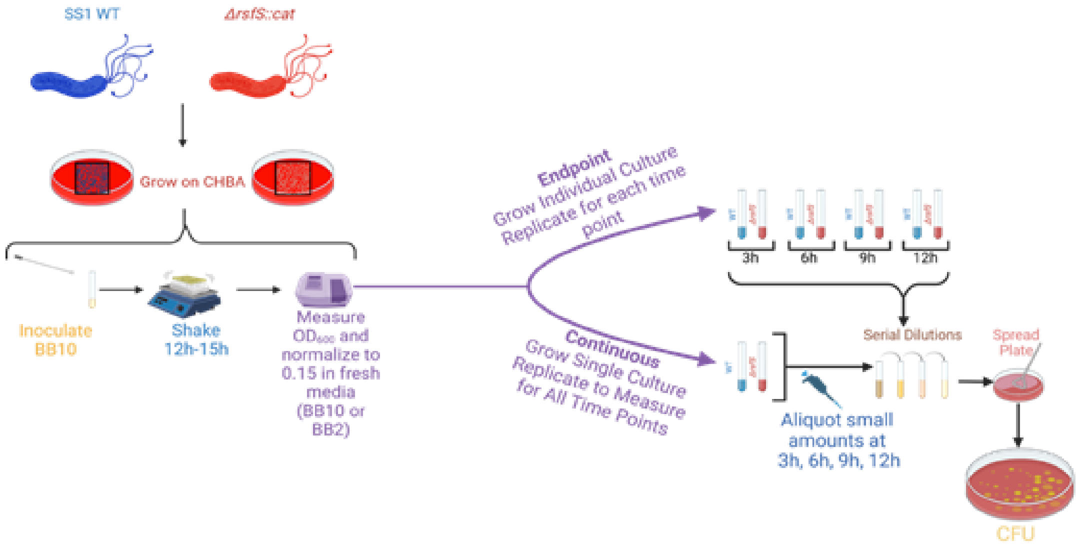


**Supplemental Figure 2. Endpoint versus Continuous Growth Curve Experiments.** Two approaches to growth curves were used to fully define potential growth defects in the Δ*rsfS* strain: endpoint and continuous growth. For the endpoint approach, multiple samples were created from the same initial starting culture, and one removed at each time point. For the continuous approach, a single initial starting culture was created and sampled repeatedly.

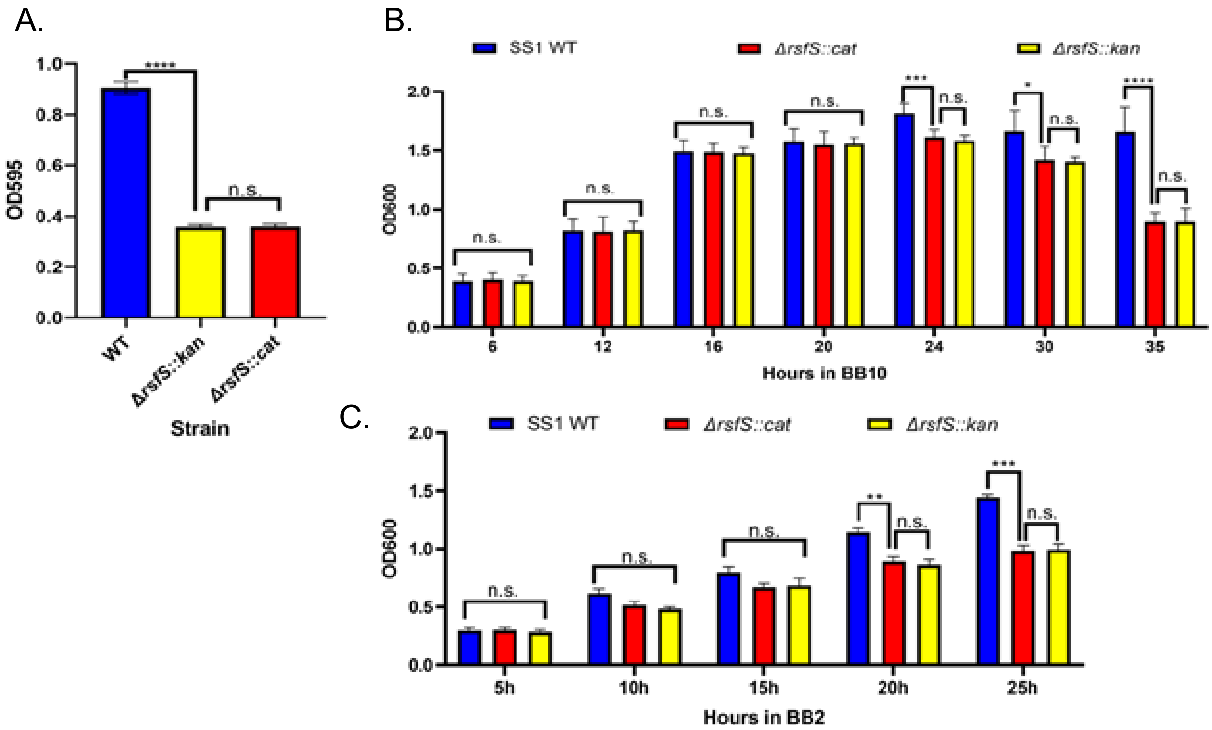


**Supplemental Figure 3. Phenotypic verification of a second Δ*rsfS* strain.** *In vitro* assays were conducted with the Δ*rsfS::kan* strain to confirm that the defective phenotypes observed in the Δ*rsfS::cat* strain are inherently due to the genetic deletion of *rsfS*. **(A)** Three biological replicates of the positive control, SS1 WT (mean= 0.905, technical replicates=71), and the negative control, Δ*rsfS::cat* (mean= 0.359, technical replicates= 70), were grown to compare to six biological replicates of the parent Δ*rsfS::kan* strain (mean=0.358, technical replicates= 152). T-test analysis with Welch's correction revealed a statistically significant defect p value <0.0001, ****, between the WT and the Δ*rsfS::kan* but no significant differences between Δ*rsfS::kan* and Δ*rsfS::cat* **(B)** Strains were grown in BB10 and statistically significant differences were not observed between WT and the Δ*rsfS::kan* until the stationary phase timepoints t= 24h, 30h and 35h with T-test analysis revealing p value <0.0001, ***,*,**** respectively but no significant differences between Δ*rsfS::kan* and Δ*rsfS::cat*. **(C)** Strains were grown in BB2 and statistically significant differences were not observed between WT and the Δ*rsfS::kan* until the later timepoints t= 20h and 25h with T-test analysis revealing p value <0.0001, **,*** respectively but no significant differences between Δ*rsfS::kan* and Δ*rsfS::cat*.

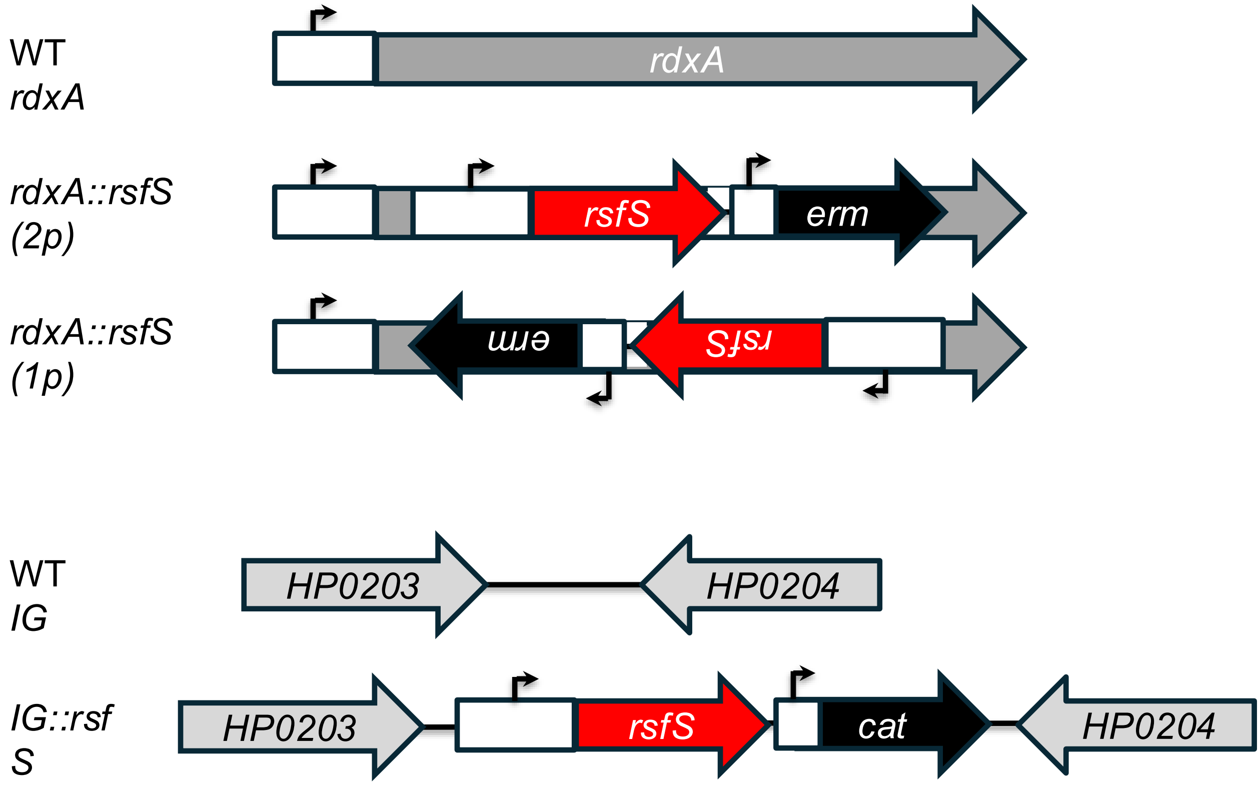


**Supplemental Figure 4. Designs for *rsfS* complementation constructs.** Plasmid inserts were designed with 200 bp upstream of the *rsfS* gene to include the predicted transcriptional start site, potential promoters (arrowheads), and any regulatory sequences. This segment along with a selectable marker, either erythromycin resistance gene (*erm*) or chloramphenicol resistance gene (*cat*) for selection and flanking with either *rdxA* or intergenic (IG) loci sequences to promote integration of the *rsfS* gene at those sites. For the *rdxA* integrated constructs, there are two versions. One has *rsfS* oriented in the same direction as *rdxA*, and thus may be transcribed from both promoters (2p). The other has *rsfS* oriented in the opposite direction as *rdxA*, and thus is predicted to be transcribed only from its own promoter (1p).

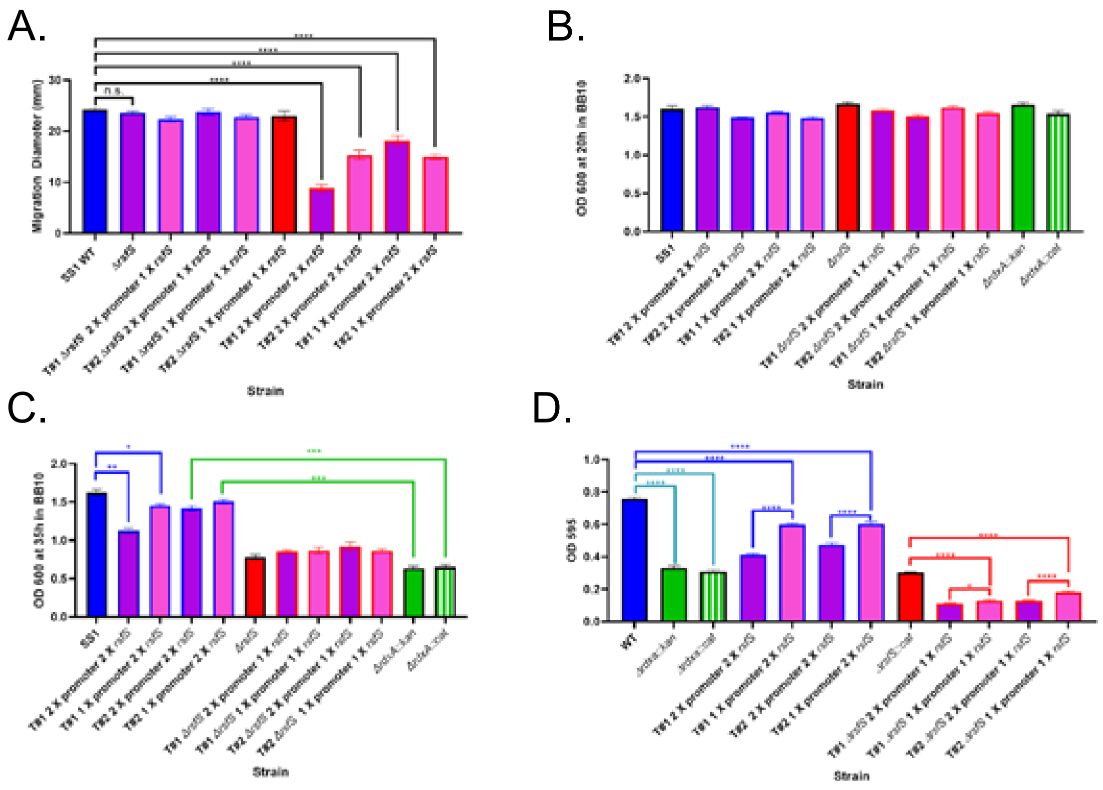


**Supplemental Figure 5. Phenotypes associated with *rsfS* expression and complementation in *rdxA*.** *rsfS* was integrated in the *rdxA* locus in either the WT or ΔrsfS background in two independent transformations, T#1 and T#2. Two variants of the *rdxA*:: *rsfS* mutants were assessed (Supplemental Fig. 5): 2P with both the rdxA and rsfS promoters, and 1P with only the *rsfS* promoter. **(A)** Soft agar assays were conducted with 12 biological replicates with 45 and 60 technical replicates respectively. Students T test was used to compare diameters to that of WT (p value <0.0001, ****). **(B)** Exponential phase growth was examined at the 20h time point, using the endpoint approach. No significant statistical difference was found across all strains relative to the SS1 WT control using Students T test. Four-five biological replicates were used. **(C)** Stationary phase $OD_{600}$ values were examined after 35h of growth. (p value <0.0001, *, **, ***). Double copy mutants were compared to WT (Blue Bracket) and Δ*rdxA* mutants (Green Bracket) because the phenotypes were partially rescued by the *rsfS* insertion despite the loss of function mutations. **(D)** Biofilms were grown in BB2 and analyzed by crystal violet staining. Three-four biological replicates per strain with 164-246 technical replicates were used. Students T test was used to compare the indicated strains (p value <0.0001, *, ****). Control strains WT and Δ*rdxA* mutants (Teal Bracket) were. WT strains were compared to double copy mutants containing single or double promoters (Blue Bracket). Δ*rsfS* mutant was compared to Δ*rsfS* Δ*rdxA::rsfS* (1 X *rsfS*) mutants containing single or double promoters (Red Bracket).
